# Leaving No Stone Unturned: Delineating the Distribution Range of the White Striped Viper-Gecko (*Hemidactylus albofasciatus*)

**DOI:** 10.1101/2025.05.03.651999

**Authors:** Prathamesh Amberkar, Siddharth Mandke

## Abstract

Delineating the distribution ranges of a species and understanding factors affecting them is not only a central theme in biogeography but is also important for their conservation. The International Union for the conservation of Nature (IUCN) Red list assess the extinction risk of the species using species range size as one of the criterion. The White-striped viper gecko (*Hemidactylus albofasciatus*) was categorized as ‘Vulnerable’ under this criterion. However the assessment was inaccurate and based on literature surveys. Moreover, the distribution range of the species is also unknown. To bridge this gap, we used occurrence data collected from opportunistic surveys and citizen science data to build ecological niche models (ENM) for the gecko. The suitable habitat were later sampled to delineate the accurate range of the species. We found, two rivers acting as geographical barriers, marking the northern and southern extent of the species’ distribution, although there was suitable habitats, for a several kilometre on either side of the barriers. We calculated the area of occupancy and extent of occurrence as 480 km^2^ and 1,609 km^2^, respectively, suggesting that the IUCN status of the species should be further elevated to ‘Endangered’.

## INTRODUCTION

Understanding the factors influencing species distribution is a central theme in biogeography. Within the same clade, species often exhibit significant disparities in their distribution ranges—some are widespread, while others are restricted to a single locality or specialized habitat (**Gaston, 1996**). These differences arise due to variations in life history traits, morphology, historical geological events, and species interactions (**Gaston, 2009**). Species with limited distribution ranges tend to have smaller population sizes and lower genetic diversity, making them more vulnerable to extinction (**Meiri et al., 2018**). Consequently, accurately delineating species ranges is crucial from a conservation perspective, as it helps prioritize resource allocation toward species at greater risk.

The International Union for the conservation of Nature (IUCN) Red list is one of the major tools for assessment of extinction risk worldwide. These assessments have been instrumental in decision making, policy implementation, funding and research **(Rodrigues et al., 2006**). To prioritize species conservation, the species are listed under various categories using five criteria, based on their extinction risks – A) population size reduction—significant decline in the population either over 10 years or three generations B) geographic range—the size and fragmentation of a species’ range C) small population size and decline—low number of mature individuals D) very small or restricted population—very small population size or restricted population regardless a trend and E) quantitative analysis—using models to estimate probability of extinction in the wild for species with limited data. Since a majority of these criteria require data of the population size, which is unavailable and difficult to procure for a majority of the species, most of these assessments are carried out using criterion B, i.e. geographic range sizes. This criterion includes two parameters: Extent of Occurrence (EOO), the area contained within the shortest continuous boundary that encompasses all known records of the species and Area of Occupancy (AOO), the area within the EOO where the species is actually found (**Gaston, 1991**). Species with an EOO of less than 20,000 sq. km are categorized as ‘Vulnerable’, those with an EOO of less than 5,000 sq. km as Endangered, and species with an EOO of less than 100 sq. km as Critically Endangered.

However, the distribution ranges of many species are unknown, referred to as the ‘Wallecean shortfall’. A couple of frameworks have been proposed to address this gap (**Farquhar et al., 2024; Muliya et al., 2021**). These frameworks revolve around constructing ecological niche models (ENM) using environmental and topographical variables, as well as collating species occurrence data from open-source databases like iNaturalist and the Global Biodiversity Information Forum (GBIF). Despite their utility, these frameworks have limitations. For some species, very few occurrence records are available due to factors such as the remoteness or inaccessibility of their habitats or their cryptic nature, making them difficult to detect. Furthermore, ENM typically predicts the fundamental niche of a species—the potential area it could occupy based on environmental suitability. However, the realized niche, the actual area where a species is found, can be much narrower **(Araújo and Guisan, 2006**). This discrepancy arises because other factors, such as geographical barriers, dispersal abilities and other biotic interaction, also influence species distribution (**Wiens, 2011**), but these are not accounted for in many modelling approaches.

The White striped viper gecko (*Hemidactylus albofasciatus*, Grandison and Soman 1963) (**Fig. 1a**) is a small-bodied, (adult, snout to vent length - 38 mm), ground-dwelling, nocturnal species confined to open habitats on the lateritic plateaus in the Konkan region of Maharashtra state, India (**Fig. 1b**). The threat assessment of the species was carried out in 2011 and the species was categorized as ‘Vulnerable’, based on the criterion ‘B1ab’, the geographic range size. The data used to assess the IUCN status of the species is based on five localities collated from literature surveys but no systematic field surveys were conducted (**IUCN, 2011**). This gecko is known from its type locality, Dorle, Ratnagiri district and a few other localities around its type locality but the exact distribution of the gecko is so far not known **(Gaikwad et al., 2009; Mirza and Sanap, 2012**). There are many regions north and the south of the type locality, with presumably suitable habitats which haven’t been sampled yet.

**Figure 1.**
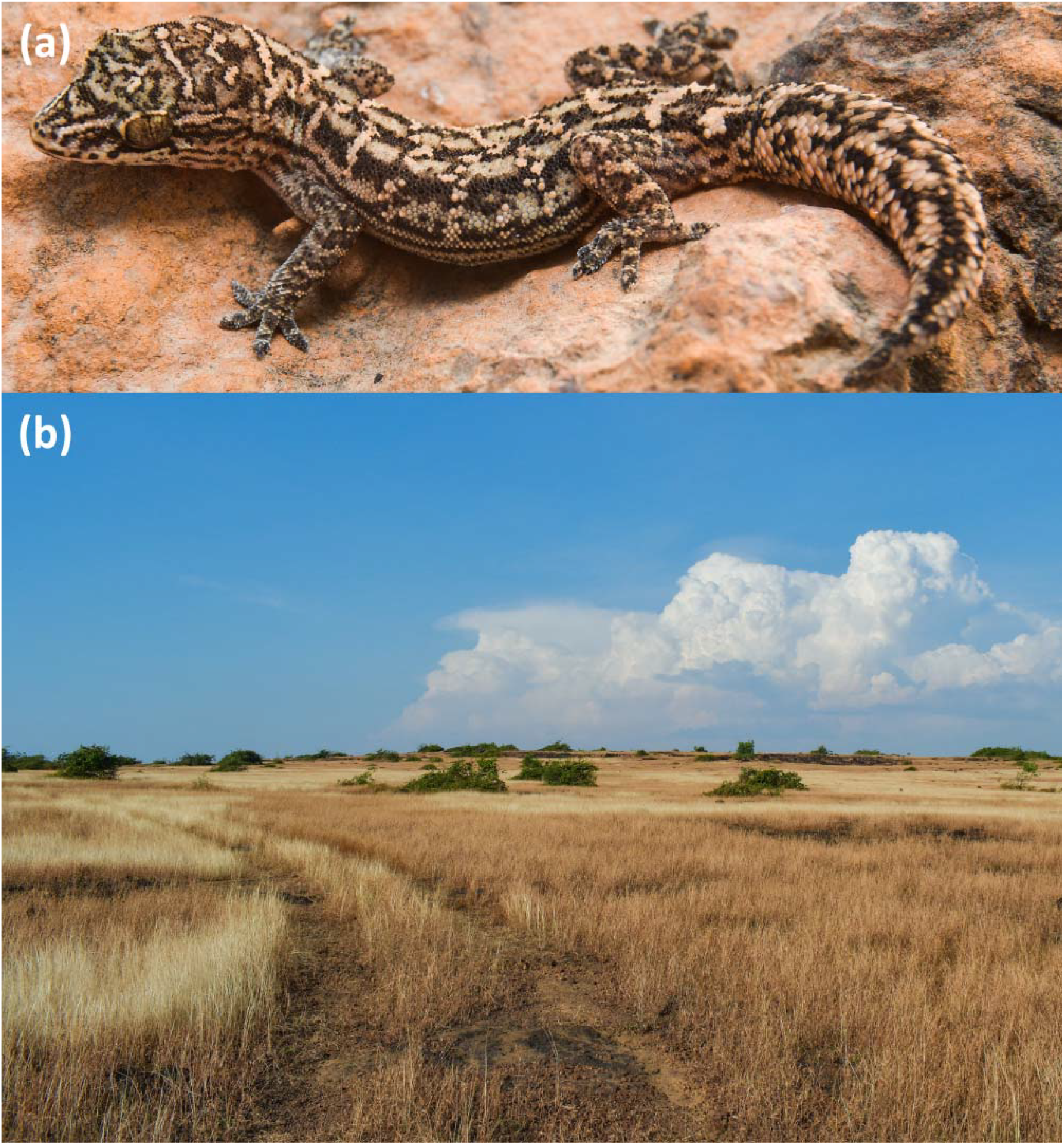
Representative pictures of the White-striped viper gecko (Hemidactylus albofasciatus) (a) and its habitat, the open lateritic plateaus (b).

In this study, we used an integrative framework to delineate the distribution range of this species, with very few occurrence records. First, we identified the suitable habitats for the species by constructing ENM. Subsequently, we conducted field surveys to sample habitats marked as suitable by the ENM to better understand the species’ distribution range. The landscape where the species is currently known—the lateritic plateaus of Konkan—is an open natural ecosystem. These plateaus are at an elevation of 50–200m asl and run parallel between the western escarpment of the Northern Western Ghats and the west coast of India in the districts of Ratnagiri and Sindhudurg, in Maharashtra, India (**Fig. 2a**). The landscape is a mosaic of open habitats, forests and human settlements (**Watve, 2013**). These plateaus are not continuous and are fragmented by several rivers flowing from the Western Ghats into the Arabian Sea, parallel to each other. Given the species’ poor dispersal ability, we hypothesize that these rivers may act as barriers to their dispersal.

**Figure 2.**
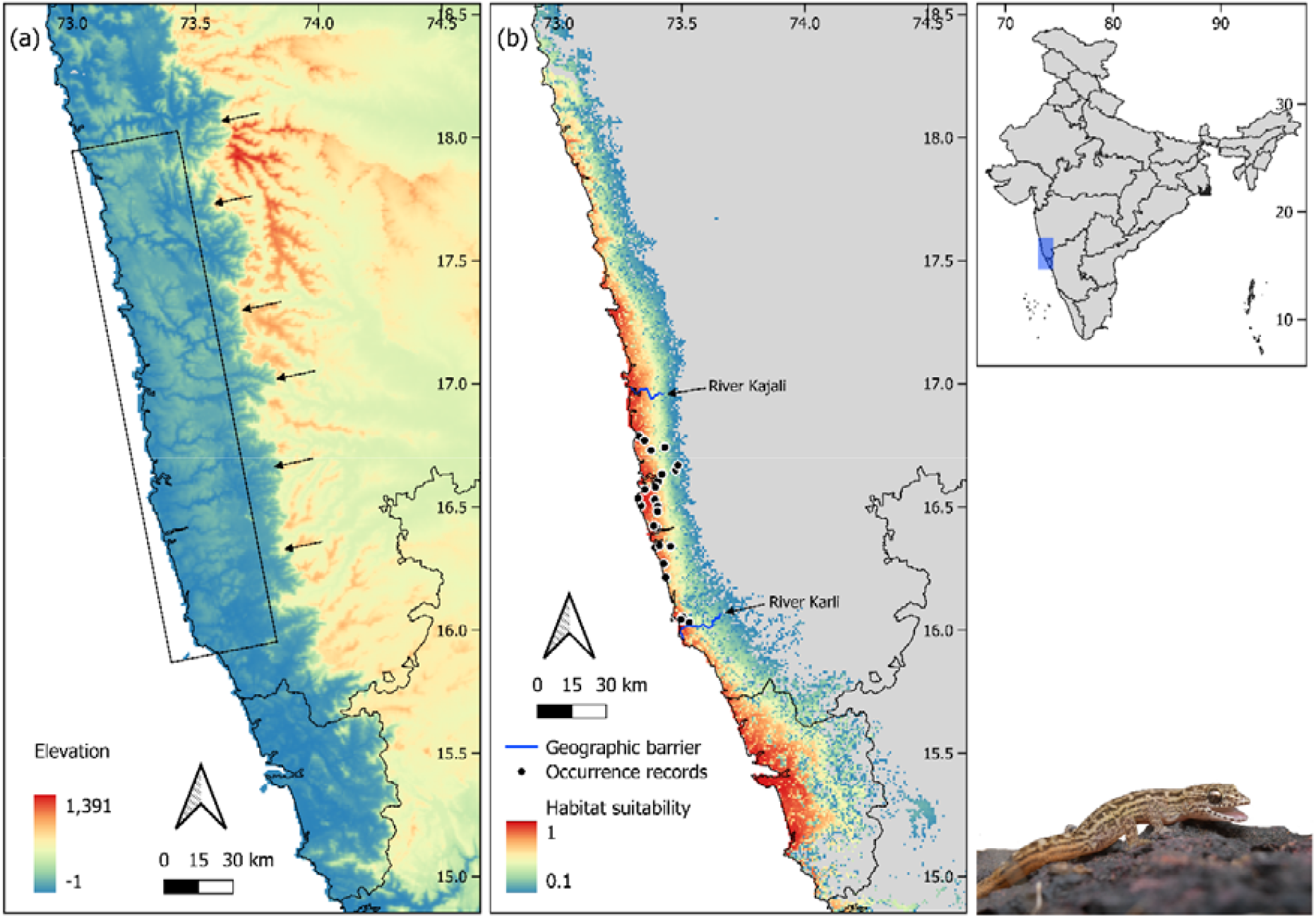
(a) Digital elevation map (elevation in meters) of the Konkan region and the Northern-Western Ghats. The western escarpment of the Northern-Western Ghats is denoted by the black arrows, showing a sharp change in elevation. The rectangle with black outline is highlighting the Konkan region. (b) Habitat suitability map of H. albofasciatus highlighting suitable habitats on either side of the rivers, where the species was absent. The black dots with white outline indicate the occurrence records used to construct ENM, collected from opportunistic surveys and open-source databases.

## METHODS

### Occurrence records of *H. albofasciatus*

We collected occurrence data for *H. albofasciatus* from opportunistic surveys we conducted in the region, supplemented by open data sources such as iNaturalist and the Global biodiversity information facility (GBIF) and published literature. Since *H. albofasciatus* is a lesser known species, it can be sometimes be mistaken as a juvenile of other commonly found *Hemidactylus* species in the region such as the Amboli brookish gecko (*H. varadgirii*). To overcome this bias while collecting iNaturalist records, we drew a boundary over the Konkan region using the ‘Explore’ option and used ‘Reptilia’ and other keyword such as ‘geckos’ and ‘lizards’ as a filters. All these search results up to August 2024 were scanned to look for records which were *H. albofasciatus* and wrongly identified as some other *Hemidactylus* species. Moreover, to protect ‘threatened’ species, iNaturalist adds uncertainty of ~ 30 km to otherwise precise occurrence records. These occurrence records, with induced bias could result in misleading ENM (**Contreras-Díaz et al., 2023**). Hence, to minimize this bias, we communicated with the observer for accurate occurrence records. We collected a total of 36 occurrence record of the species. When multiple occurrence records were within 1 km^2^ radius of each other, only one was retained to ensure that each cell in the predictor layers contained no more than one presence record. Finally, 25 occurrence records were retained and used for further analysis.

### Environmental variables

The species is found in open habitats, with scattered shrubs with no tree cover. Hence, we used tree-cover as one of the variables for my ENM. This layer was obtained from Hansen et al. 2013, where tree-cover is defined as canopy closure for all vegetation taller than 5m in height encoded as a percentage per output grid cell, for the year 2000 (**Hansen et al., 2013**). Since the plateaus, where the species is found are at an elevation of 50–200m asl, we used topographical layers such as digital elevation maps (DEM), aspect and slope. Further, we collected raster datasets of bioclimatic variables from WorldClim (**Fick and Hijmans, 2017**). Climatic variables such as mean temperatures of the wettest and driest quarters (BIO8 and BIO9) and mean precipitation of the warmest and coldest quarters (BIO18 and BIO19) were not considered for analyses as they potentially leave spatial artefacts in the data (**Campbell et al., 2015**). Moreover, since the species is nocturnal environmental variables such as BIO2— mean diurnal range and BIO3—isothermality were also excluded from the analysis. The spatial extent of these environmental layers were trimmed to only include the Konkan-Malabar region and the Western Ghats. To account for the spatial resolution between tree-cover and other environmental and topographical layers these layers were resampled to 1 sq. km. using bilinear interpolation, using the function ‘*resample*’ in the R package ‘*raster*’. All the layers were tested for collinearity and the Pearson’s correlation coefficient (|*r*|>0.75) (**Dormann et al., 2013**) between variables pairs was used to identify strongly correlated variables. Finally, 10 weakly correlated variables were retained for further analyses. These include Tree-cover, Aspect, DEM, Slope, BIO1 (Annual mean temperature), BIO4 (Temperature seasonality), BIO12 (Annual precipitation), BIO14 (Precipitation of driest month) and BIO15 (Precipitation seasonality) (**Fig S1**). Although BIO1 and DEM were highly negatively correlated, we retained both these predictors since they both were important for the species.

### Ecological Niche Model

The aim of the ENM was to earmark the suitable habitats for the species for further field sampling. Hence, considering the less number of occurrence data points for the species, we constructed the model using MaxEnt. MaxEnt incorporates presence-only data and environmental variables, and perform better compared to other algorithm while working with a few occurrence data points (**Phillips et al., 2006**). Optimal MaxEnt settings were obtained using ‘*ENMeval*’ package in RStudio (**Kass et al., 2023**). This package runs MaxEnt across various combinations of feature classes and values of regularization multiplier to enable comparisons of model performance. These models were built with regularization multiplier options from 1–5 and six different feature class combination – Linear, Linear + Quadratic, Hinge, Linear + Quadratic + Hinge, Linear + Quadratic + Hinge + Product, Linear + Quadratic + Hinge + Product + Threshold (Hence 30 candidate models).

To prevent over-fitting and limit complexity, MaxEnt uses regularization. The package, *ENMeval* suggested a regularization multiplier of 2.0 and linear function (L) as a feature class (**Table S1**). Since MaxEnt uses presence-only data, its uses pseudo-absence points or background points which are comparatively high compared to the presence points. MaxEnt assumes that the entire area of interest has been systematically sampled. But the occurrence data was compiled from open data sources and published literature where sampling efforts were not uniform across the study area, leaving a spatial bias in sampling. This spatial bias could lead to inaccurate model because of over-representation of certain environmental features of the more accessible and extensively surveyed areas. To overcome this spatial bias, we used a bias file to preferentially select background points from areas with more sampling efforts, and helps the model avoid overfitting to regions with more occurrence data **(Kramer □ Schadt et al., 2013; Phillips et al., 2009**).

We generated 20 replicates of each model by bootstrapping to estimate the variability and used 10,000 background points. The jackknife test in MaxEnt was used to estimate the contribution of each variable to the final model, both, when the variable is used in isolation and when is left out from the other set of predictors. MaxEnt generates average suitability maps (of the 20 replicates). This map highlights suitable habitats with an estimated probability of the presence of the species for the desired geographic extent, with values ranging from 0 (unsuitable) to 1 (suitable). Moreover, MaxEnt also generates response curves for each environmental variable chosen (**Phillips et al., 2006**).

### Evaluating the model performance

To evaluate the performance of the ENM, we used the area under the curve (AUC) of the receiver operating characteristic curve (ROC), a widely used threshold-independent metric. The AUC is obtained by plotting sensitivity (the proportion of correctly predicted presences) against 1–specificity (the proportion of incorrectly predicted absences). An AUC value greater than 0.9 was used as an indicator of robust model performance. However, the use of AUC for presence-only models has been criticized, particularly due to low sampling prevalence—the relatively small number of presences compared to background points. When a large number of background points are environmentally distant from known presences, specificity tends to increase artificially, leading to inflated AUC values. To address this potential bias, a bias file was used during background point selection, ensuring that background points were drawn from areas with comparable sampling effort or environmental conditions (refer to the section—Ecological Niche Model). This approach reduces the risk of artificially high AUC values.

### Sampling for *H. albofasciatus*

*H. albofasciatus* is found only in open habitats on the plateaus (**Gaikwad et al., 2009**). Hence, we used an estimated probability of > 0.3 as a threshold to choose sampling locations since the model highlighted areas towards forested areas of the foothills of the Western Ghats. Grids of 500 m x 500 m were laid over the suitable habitats for systematic sampling. The landscape is a mosaic of various land-uses such as mango orchards, cashew plantation, paddy, villages, rock quarries and other built-up areas. The species is scarce in as orchards and paddy fields, and prevalent in open habitats. Hence, we surveyed grids which were overlapping with open habitat with no forests or land-uses changes. The species is nocturnal and ground-dwelling and actively forage at night, making it difficult to spot. They seek refuge under rocks during the day, making it easier spot them by flipping rocks. Hence, we carried out surveys between 0900 hr – 1800 hr by actively looking for them under rock with one other observer. The EOO and AOO were calculated using the package ‘*BiodiversityR*’ in the Rstudio (**Kindt, 2025**). The EOO was represented using a convex hull, constructed using QGIS (version - 3.38.3).

## RESULTS

### Ecological niche model

The average habitat suitability (20 replicates) map of the species suggested areas between 17°N and 15°N and the foothills of the Western Ghats marking the eastern extent of the species (**Fig.2b**). The optimal model performed well with an AUC of 0.994. The predictors BIO4 (49%), BIO15 (19.5%) and DEM (12.8) were important predictors of habitat suitability. The topographic variables, slope and aspect, and the environmental variable, BIO1 contributed relatively less to the model. The response curves for each predictors from the MaxEnt model and the results of the jackknife are provided in the Supplementary materials.

### Distribution range of *H. albofasciatus*

We sampled a total of 511 grids, each of 500 x 500 m. Of these total grids sampled, the species was present in 124 grids and not encountered in 387 grids (**Figure S2**). We encountered over 219 individuals of the species in the Ratnagiri and Sindhudurg districts of Maharashtra. Further, the EOO and AOO of the species was calculated to be 1,609 km^2^ and 480 km^2^, respectively.

As hypothesized, we found geographical barrier (rivers in this case) shaping the distribution of the species. River Kajali in the Ratnagiri district (**Fig. 3b**) marked the northern boundary of the species. Individuals of the species were abundant a few hundred meters towards the south of the Kajali River where there was suitable habitat highlighted in the habitat suitability map. Towards the north of the river, the species was absent even though the ecological niche model suggested suitable habitats there. Similarly, towards the south, in Sindhudurg district, river Karli (**Fig.3b**) marks the southern boundary to the distribution of the species and barrier to their dispersal. The species are abundant towards the north of the river Karli where there is suitable habitat but absent towards south. The species was present along all the plateaus between the rivers except for a few isolated plateaus towards the south.

**Figure 3.**
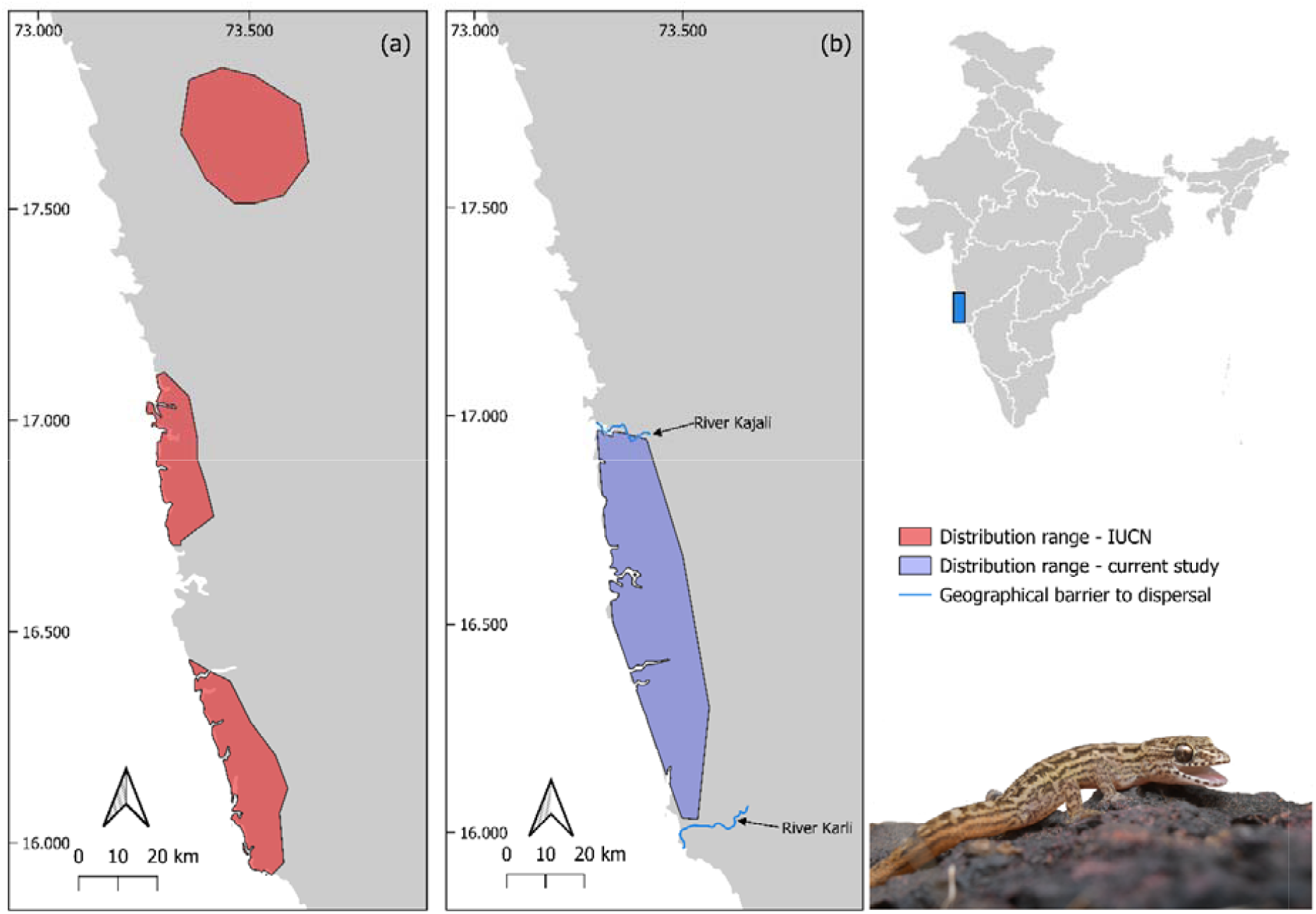
The distribution range map of H. albofasciatus from the present study (b) compared to the distribution map of the IUCN (a).

## DISCUSSION

For conservation of a species it is first important to delineate the distribution of the species, threatened with extinction. This helps policy makers in prioritizing areas for conservation. The White-striped viper gecko is such a species with a hitherto unknown geographic range. Hence, we first build ENM and then sampled the suitable habitats to accurately to delineate the distribution range of the species. We found two river acting as a geographical barrier to the dispersal of the species, marking the northern and southern extent of their distribution.

### Factors affecting the distribution of *H. albofasciatus*

Understanding factors that shape the spatial limits of a species are one of the central themes in biogeography. Many factors such as environmental variables, geographical barriers, dispersal abilities and biotic interaction (both negative-interspecific competition between species and positive – mutualism (**Narayanan and Shaw, 2024**)) can result in absence of a species from suitable habitats (**Wiens, 2011**). In the case of *H. albofasciatus*, the distribution range was restricted between two rivers, Kajali in the north and Karli in the south, although the ENM highlighted suitable habitat on either sides of these rivers (**Fig. 2b**). This suggests that these rivers marked the northern and southern boundary of the distribution of *H. albofasciatus*. Although several rivers flow parallel from the Western Ghats into the Arabian Sea, between the Kajali and Karli rivers, the species distribution is restricted by these two rivers. Why do these particular rivers define the species’ distribution, while others do not act as barriers? The escarpment of the Western Ghats was formed due to the rift-flank uplift as a result of rifting and separation of the Seychelles (~65 mya). The Escarpment was later eroded and by the several west flowing rivers and recced further inland, away from the coast. The eroded surface are the current costal lateritic plateaus, which were later fragmented by these rivers **(Kale, 2009; Radhakrishna et al., 2019; Radhakrishna and Joseph, 2012; Widdowson and Cox, 1996**). It is likely that *H. albofasciatus* once occupied a much broader range, extending from the coastal lateritic plains to the plateaus further east, in the Western Ghats. However, the formation of river systems and the establishment of dense forests along the Western Ghats would have created dispersal barriers, leading to population isolation. One such isolated lineage may have undergone allopatric speciation, giving rise to *Hemidactylus sataraensis*—a small-bodied, ground-dwelling gecko, sister to *H. albofasciatus* now restricted to open lateritic plateaus within the Western Ghats, several kilometres east of the coastal plains. With the fragmentation of the landscape, *H. albofasciatus* likely became restricted to the coastal lateritic plains, while *H. sataraensis* persisted on the elevated plateaus of the Ghats.

### IUCN status of *H. albofasciatus*

Although the IUCN Red List is a crucial tool for conservation, many researchers have raised concerns about its risk assessment methodology, highlighting flaws in the evaluation of numerous species (**Caetano et al., 2022; Edgar, 2025; Palacio et al., 2023; Seminoff and Shanker, 2008; Webb, 2008**). The risk assessment of *H. albofasciatus* was carried out in 2011, based solely on literature surveys, without systematic field surveys. The species’ range was represented using a convex hull polygon, which resulted in three distinct polygons, highlighting three different populations (**Fig. 2a**) (**IUCN, 2011**). Contrastingly, the convex hull polygon constructed from the occurrence point of the current study resulted in a single continuous polygon (**Fig. 2b**). Moreover, the representation of the species’ distribution range, constructed using occurrence data from published literature is inaccurate. While the two southern polygons contain confirmed occurrence records, the northernmost polygon lacks any supporting occurrence data, making its inclusion questionable (**IUCN, 2011**). While applying criterion B, the general threshold either on EOO and AOO must be first met. Later, the taxon must meet at least two of the three options listed for criterion B – (a) severely fragmented distribution (b) continuing decline and (c) extreme fluctuation (**IUCN, 2024**). *H. albofasciatus* was earlier categorized ‘Vulnerable’ under the criteria ‘B1a’ and no data on sub criterion ‘a’.

In the present study, the EOO and AOO were calculated as 1,609 km^2^ and 480 km^2^ respectively. Although the species meet the threshold of the EOO and AOO for reclassifying it as ‘Endangered’ under the criterion B, the data required to classify it further, in the latter categories is not available. But, considering that the habitat that the species is found, the open lateritic plateaus, experience a rapid change in anthropogenic land-use patterns, which are negatively affecting the species (**Amberkar and Mungikar, 2024; Jithin et al., 2023**), the species must be categorized as ‘Endangered’.

Among the various criteria used to assess extinction risk for reptiles, Criterion B— geographic range size—is the most commonly applied (**Meiri et al., 2023**). This criterion is quantified using two key measures, the AOO and the EOO. However, both measures have certain limitations. EOO is typically calculated as the convex hull polygon encompassing all known occurrence points (**Gaston, 1991**), often leading to an overestimation of range size by including large areas where the species is absent. This limitation is particularly relevant for species with fragmented distributions and species confined to only a particular habitat within the landscape, as the method does not account for habitat suitability or ecological constraints. Additionally, EOO is sensitive to sampling biases—if occurrence records are sparse or biased toward accessible locations, a few outlying records can disproportionately inflate the estimate. The EOO for *H. albofasciatus*, was calculated to be 1,609 km^2^. EOO calculations may overestimate its range by including unsuitable forested habitats adjacent to the lateritic plateaus where the species is not found. Moreover, there were a few plateaus towards the south of the distribution of the species, where the species was absent but still were overlapping with the EOO. Thus, while EOO provides a broad geographic context, it should be interpreted alongside finer-scale measures such as the AOO.

The AOO is highly sensitive to the spatial scale (grain size) at which it is measured. Smaller grain sizes yield lower AOO estimates, necessitating greater sampling effort to obtain an accurate value. Consequently, to achieve a reliable AOO estimation, it is recommended to comprehensively map the species across its entire potential range (**Marsh et al., 2023)**. If there are unsampled areas where the species is present but unrecorded, AOO will be underestimated, making it particularly vulnerable to incomplete data. A standard grain size of 2×2 km as recommended by the IUCN. Using this method, the AOO of *H. albofasciatus* was calculated to be 480 km^2^, calculated as the sum of occupied grid cells containing known occurrence points. However, as discussed, this estimate may vary since some suitable grid cells within the lateritic plateaus could remain unsampled, potentially leading to an underestimation of the true AOO.

### Conservation of *H. albofasciatus*

The habitat where the species is found, the open lateritic plateaus, are classified as ‘Wasteland’ (Government of India and Department of land resources, 2019) due to their barren appearance (**Watve, 2013**). These plateaus are subjected to various land-uses such as mango orchards, paddy and stone quarries which negatively affect the wildlife inhabiting these plateaus, including *H. albofasciatus*. This species is more prevalent in unaltered open plateaus compared to the other land uses. Notably, none of its known distribution falls within any protected area. Since the distribution of the species is known, designating a few plateaus as Biodiversity Heritages sites, recognized under the Biological Diversity Act, 2002 of the Government of India, could be an effective conservation strategy. This will help ensure that this gecko and other species inhabiting the plateaus are conserved and at the same time, the local communities are not restricted by using the natural resources found on the plateaus (**The Biological Diversity Act, 2002**).

With rising global temperatures due to climate change, species are often forced to shift their ranges to more suitable habitats (**Thomas, 2010**). However, *H. albofasciatus* has a highly restricted range, low dispersal ability, and is confined to open lateritic plateaus. Given the rapid pace of climate change, the species may struggle to track suitable habitats, increasing its risk of extinction.

### Future research questions

The distribution of *H. albofasciatus* appears to be influenced by geographical barriers, particularly rivers. Given that multiple rivers originating from the Western Ghats flow parallel to each other, fragmenting the plateaus, it is plausible that these rivers similarly fragment the populations of *H. albofasciatus*. If these rivers have indeed played a role in shaping the population structure of the species, we might expect to find distinct, independently evolving lineages on each isolated plateau. Investigating genetic divergence among populations separated by these rivers could provide insights into historical gene flow, vicariance, and the role of rivers as isolating barriers in shaping the evolutionary history of the species.

While abiotic factors play a crucial role in determining species distributions, biotic interactions—both competitive and predatory—can further shape their distribution **(Wiens, 2011**). Across ecosystems, such interactions are known to operate at multiple spatial scales, influencing species assemblages and habitat preferences. In the case of *H. albofasciatus*, competitive exclusion and predator-prey dynamics may be key factors restricting its occurrence to the open lateritic plateaus. On these plateaus, *H. albofasciatus* is often found sheltering under rocks, coexisting with other lizard species such as *Ophisops jerdonii* and *O. beddomei*. Despite sharing the same feeding guild, these species are primarily diurnal, which may facilitate their coexistence with the nocturnal *H. albofasciatus* through temporal niche partitioning. However, in the adjacent forested habitats, the presence of generalist gecko species that exploit similar resources could lead to strong interspecific competition. If these generalists outcompete *H. albofasciatus* for food or refuge, this could explain the gecko’s confinement to open habitats, where such competition is reduced. Moreover, there are various other species of snakes found in forests adjacent to the open habitats. These snakes could be potential predators to the species further restricting their range. Additionally, the forests surrounding these plateaus harbor various snake species such as the Buff-striped Keelback (*Amphiesma stolatum*), Wolf snakes (*Lycodon sp*.), Kukris (*Oligodon sp*.) etc which could be potential predators of *H. albofasciatus*, further restricting its range. The combined pressures of predation and interspecific competition may reinforce the gecko’s preference for open habitats, where both threats are reduced.

## DATA AVAILABILITY STATEMENT

The occurrence data of the species collected during the field surveys and the R scripts used to derive optimal settings while using MaxEnt will be deposited in an open source data repository once the article has been published.

## ACKNOWLEDGEMENT

We would like to thank Mr Aritra Biswas, Mr Chaitanya R and Dr Aparna Lajmi for their valuable comments on the manuscript. PA would also like to thanks Ms. Christi Sylvia for her insightful comments on the study design.

