## Supplementary material for "Leaving No Stone Unturned: Delineating the Distribution Range of the White Striped Viper-Gecko (*Hemidactylus albofasciatus*)": Figures S1 and S2

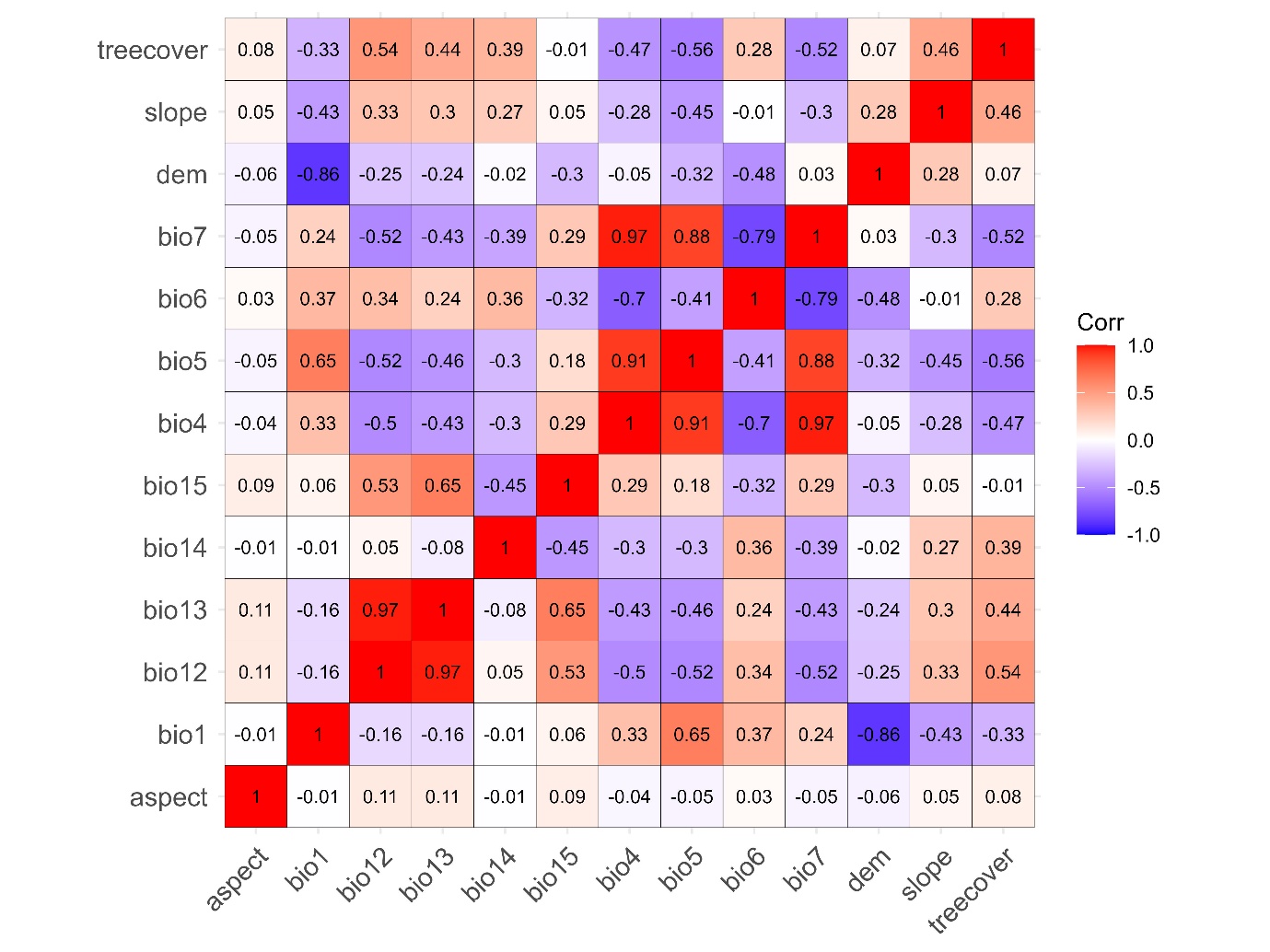

**Figure S1:** Correlation matrix of environmental and topographical variables used to shortlist variables to construct the species distribution model. Out of the 13 variables we discarded highly correlated variables (|r|>0.75) and retained 10 variables.

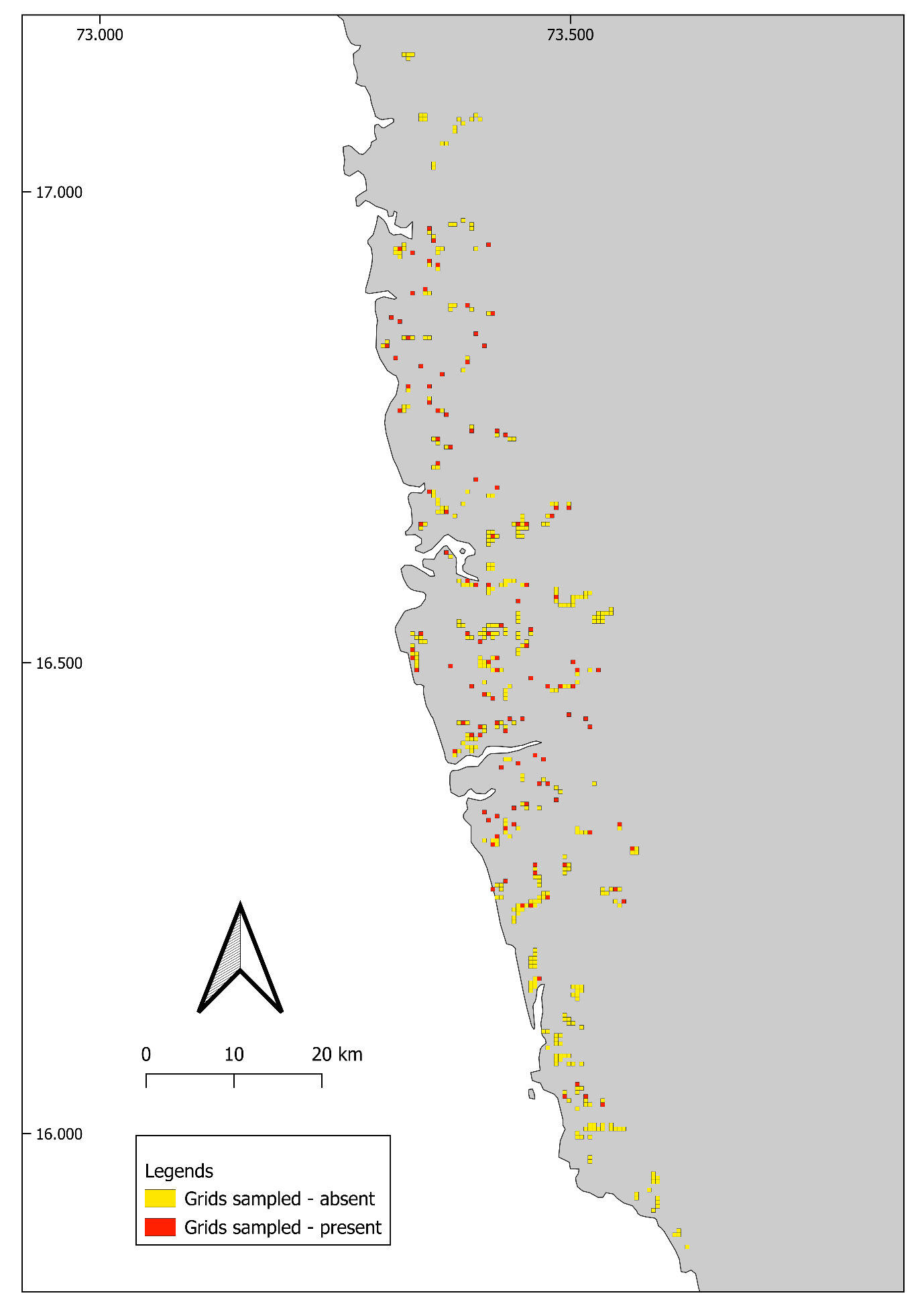

**Figure S2:** The yellow and red boxes represent grids that were sampled and the species was absent and the grids that were sampled and the species was not encountered, respectively. A total of 511 grids of 500 x 500 m each were sampled. The species was encountered in 124 grids and was not encountered in 387 grids.
